# A peptide-mediated, multilateral molecular dialogue for the coordination of pollen wall formation

**DOI:** 10.1101/2022.01.26.477813

**Authors:** Jekaterina Truskina, Stefanie Brück, Annick Stintzi, Sophy Boeuf, Nicolas M. Doll, Satoshi Fujita, Niko Geldner, Andreas Schaller, Gwyneth C. Ingram

## Abstract

The surface of pollen grains is reinforced by pollen wall components produced non-cell autonomously by tapetum cells that surround developing pollen within the male floral organ, the anther. Here we show that tapetum activity is regulated by the GASSHO (GSO) receptor-like kinase pathway, controlled by two sulfated peptides, CASPARIAN STRIP INTEGRITY FACTOR 3 (CIF3) and CIF4, the precursors of which are expressed in the tapetum itself. Coordination of tapetum activity with pollen grain development depends on the action of subtilases, including AtSBT5.4, which are produced stage-specifically by developing pollen grains. Tapetum-derived CIF precursors are processed by subtilases, triggering GSO-dependent tapetum activation. We show that the GSO receptors act from the middle layer, a tissue surrounding the tapetum and developing pollen. Three concentrically organized cell types therefore cooperate to coordinate pollen wall deposition through a multilateral molecular dialog.

**Significance Statement:** Pollen viability depends on a tough external barrier called the pollen wall. Pollen wall components are produced by tapetum cells which surround developing pollen grains within the anther. Precise coordination of tapetum activity with pollen grain development is required to ensure effective pollen wall formation. Here we reveal that this is achieved through a multidirectional dialogue involving three distinct cell types. We show that peptide precursors from the tapetum are activated by proteases produced stage-specifically in developing pollen grains. Unexpectedly, we found that activated peptides are perceived not in the tapetum, but in the middle layer, which encloses the developing tapetum and pollen grains, revealing an unsuspected role for this enigmatic cell layer in the control of tapetum development.

## Introduction

Pollen grains, housing dispersible male gametophytes, form a critical element in plant sexual reproduction. The pollen from a single plant can be carried by wind or pollinators towards multiple other individuals allowing efficient exchange of genetic material. However, the release of pollen grains into the terrestrial environment requires protection from dehydration, sunlight and other environmental stresses. Protection is provided by a multilayered pollen wall with sporopollenin, one of the most resistant biological polymers known, as the main constituent of the outermost layer (the exine) (*1*).

Pollen is produced in the anthers from diploid precursor cells, the pollen mother cells (PMC). Each PMC undergoes meiosis forming four haploid cells. These are temporarily held together as tetrads before they are released as individual immature microspores (*2*, *3*). The subsequent assembly of the pollen wall is a highly dynamic multistep process involving the sporophytic tissues that surround developing pollen grains. In Arabidopsis, these comprise four concentric layers of cells: the tapetum (most internal cell layer), middle layer, endothecium and epidermis (outer cell layer) (Fig. 1A). The tapetum contributes the most to pollen wall formation, by producing the biochemical precursors of sporopollenin and of other wall components (*1*, *3*, *4*). However, the developing pollen grains are not in direct contact with the tapetum but are suspended a matrix of largely unknown composition (often called the locular fluid), which is presumably also produced by the tapetum (*5*). The tapetum cells export pollen wall components, which must then traverse this matrix for deposition on the pollen surface to form an intact and functional pollen wall (Fig. 1B).

**Fig. 1.**
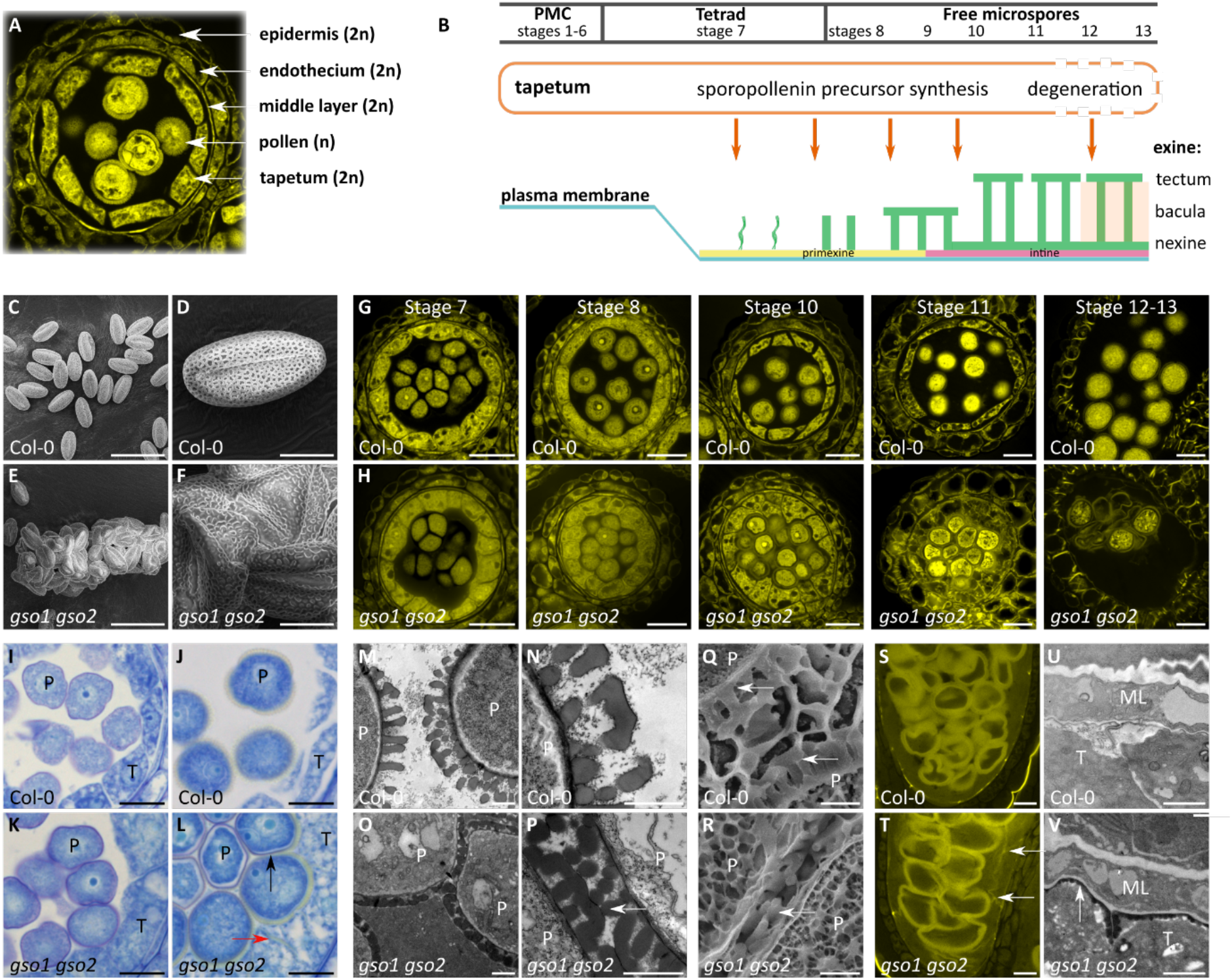
GSO1 and GSO2 receptor-like kinases are required for pollen wall formation and tapetum development. **(A)** Inside the anther the haploid (n) pollen are surrounded by four diploid (2n) sporophytic cell layers: tapetum, middle layer, endothecium and epidermis. **(B)** At the end of the tetrad stage (stage7) the tapetum cells start to synthesize and release sporopollenin precursors into the locular matrix where they ultimately attach to the surface of the pollen grains. When pollen wall construction is finished (stage 11), the tapetum cells degrade and the resulting cellular debris also adhere to the pollen surface. Pollen wall structural components are indicated: The exine is composed of nexine, bacula and tectum. PMC: pollen mother cell. **(C-F)** Scanning electron microscopy (SEM) images of the wild-type (C-D) and the *gso1-1 gso2-1* (E-F) mature pollen. **(G-H)** Pollen and anther development in wild-type anthers (G) and *gso1-1 gso2-1* anthers (H) from the tetrad stage (stage 7) to pollen release (stage 13). **(I-L)** Pollen wall formation in wild-type (I-J) and the *gso1-1 gso2-1* mutant (K-L) anthers shown by toluidine blue staining at stage 8 (I, K) and stage 10 (J, L). Black arrow indicates reddish purple staining on the surface of the pollen grains in the *gso1-1 gso2-1* mutant; red arrow indicates ectopic yellow staining between tapetum cells in the *gso1-1 gso2-1* mutant. **(M-P)** Transmission electron microscopy (TEM) of the pollen wall in wild-type (M-N) and *gso1-1 gso2-1* (O-P) pollen. Arrow indicates fused pollen wall in the mutant. **(Q-R)** Cryo-SEM of the pollen wall in wild-type (Q) and *gso1-1 gso2-1* (R) pollen. Arrows indicate the pollen walls. **(S-T)** Pollen wall material stained with Auramine-O in wild-type (S) and *gso1-1 gso2-1* (T) anthers. Arrows indicate ectopic staining in the mutant. **(U-V)** TEM showing ectopic deposition of sporopollenin-like material around the tapetum cells in the *gso1-1 gso2-1* mutant (V) (arrow) compared to the wild-type (U). Scale bars: C, E 50 μm; G-H 20 μm; D, F, I-L, S-T 10 μm; M-R, U-V 1 μm. P = pollen, T = tapetum, ML = middle layer.

The export of pollen wall components from the tapetum is under developmental control. It terminates with programmed cell death of the degenerating tapetum, liberating the final components of the pollen wall. Although the critical contribution of the tapetum to pollen wall development is uncontested, the question of whether, and if so how it is coordinated with pollen grain development, as suggested in the literature (*6*, *7*), remains unanswered.

Two extracellular structural barriers have recently been shown to be developmentally monitored to ensure physical integrity prior to their functional deployment. These are the embryonic cuticle (necessary to prevent water loss from the seedling surface at germination), and the Casparian strip (necessary for the regulation of water and ion homeostasis in roots). The integrity of both the nascent Casparian strip and the nascent embryonic cuticle, depends on intercellular signaling pathways involving GASSHO (GSO) receptor-like kinases (RLKs) and CASPARIAN STRIP INTEGRITY FACTOR (CIF)-related sulfated peptide ligands (*8*–*10*). In both cases, the diffusion of post-translationally processed ligands across the barrier into tissues containing functional receptor complexes signals defects in barrier integrity and triggers gap-filling responses.

Here we show that tapetum function necessary for the formation of a third extracellular structure, the pollen wall, is controlled through a molecular dialogue between the middle layer, the tapetum, and the developing pollen grains. This dialogue involves two previously uncharacterized CIF peptides (CIF3 and CIF4) and their cognate receptors GSO1 and GSO2.

## Results

### The receptors GSO1 and GSO2 and the sulfated peptides CIF3 and CIF4 are necessary for normal tapetum development and pollen wall formation

We noticed in previous crossing experiments that pollen from *gso1 gso2* double mutant flowers form large clumps. Scanning Electron Microscopy confirmed that in contrast to wild-type mutant pollen, which is released as individual grains (Figure 1C and D), *gso1-1 gso2-1* double mutant pollen tends to form large fused masses of misshapen grains (Figure 1E and F) with some free non-fused pollen grains remaining. This phenotype was confirmed in the independent *gso1-2 gso2-2* double mutant carrying alternative T-DNA insertions, and was rescued by complementation with the *GSO1* wild-type sequence (Figure S1A-B). The single *gso1* and *gso2* mutants produced normal pollen (Figure S1C-D).

Five ligands of GSO1 and/or GSO1 and GSO2 have already been identified; TWISTED SEED1 (TWS1) involved in embryonic cuticle formation (*9*), CIF1 and CIF2 involved in Casparian strip formation (*8*, *10*) and CIF3 and CIF4 (*11*). While pollen of *tws1* and *cif1 cif2* double mutants was unaffected, simultaneous loss of *CIF3* and *CIF4* function phenocopied the *gso1 gso2* pollen phenotype (Figure S2A-B). Occasional pollen adhesion was also observed for single *cif4* but not for the *cif3* single mutant (Figure S1E, F, M, N), suggesting that these genes do not act fully redundantly. Because CIF3 and CIF4 belong to a family of sulfo-peptides that rely on post-translational tyrosine sulfation for GSO binding and activity (*8*, *9*, *11*), we suspected involvement of the unique Arabidopsis tyrosine protein sulfotransferase (TPST). Similar pollen defects were indeed observed in the *tpst* mutant, although, as reported in the seed, the phenotype tended to be less severe than that caused by loss of ligand or receptor function (Figure S2C-D). Despite their dramatic phenotype, the *gso1-1 gso2-1*, *cif3 cif4*, and *tpst* mutants were not sterile, as viable pollen was detected using Alexander staining (Figure S3A-H) and pollen located at the periphery of the clusters was still able to germinate (Figure S3I-L). Furthermore, plants left to self-fertilize produced viable seeds, although seed number per silique was reduced compared to wild type (Figure S3M).

We traced the origin of the pollen defects by histological analyses. Anther development appeared normal in all backgrounds until stage 8 (staging according to (*12*)) the point of microspore release (see Figure 1B). From this stage onwards pollen grains were uniformly distributed within the locular matrix of wild-type anthers. Tapetal cells were well-defined and thinned progressively until they underwent cell death, prior to the release of pollen grains between stages 12 and 13 (Figure 1G; (*13*)). In contrast, in *gso1 gso2*, *cif3 cif4* and *tpst* mutants the apparent volume of locular matrix surrounding the developing pollen grains was dramatically reduced. The swollen and hypertrophied tapetum occupied a much larger volume resulting in an apparent “crowding” of the developing pollen grains and a lack of locular fluid (Figure 1H, Figure S2). Further aspects of this phenotype were related to the deposition of pollen wall components. In wild-type stage 8 anthers, the pollen wall stains reddish purple with toluidine blue (Figure 1I)(*14*, *15*). The reddish-purple staining is masked in subsequent stages by progressive deposition of the exine, which is visible in toluidine blue-stained sections as a yellow halo around developing pollen grains (Figure 1J). This is also observed for peripheral pollen grains of *gso1 gso2*, *cif3 cif4* and *tpst* mutants. However, the walls of more internal pollen grains continue to stain reddish purple, suggesting abnormal/reduced exine deposition (Figure 1 K-L and Figure S4A-D).

Both cryo-scanning (SEM) and transmission electron microcopy (TEM) analyses of pollen walls showed that many pollen grains were physically fused in the mutant backgrounds via a shared exine (Figure 1 O-P, R; Figure S4E-L), whereas the exine layers of neighboring pollen grains were clearly separated by locular matrix in wild-type anthers (Figure 1 M-N, Q). Furthermore, the normal exine structure of bacula (pillars) and tecta (connecting the apices of the bacula on the outer surface of the pollen grain) was difficult to discern in mutant pollen grains, particularly those not in direct contact with the tapetum. The bacula were often shorter in mutants (Figure 1M-R; Figure S4E-L).

In wild-type anthers, in addition to appearing yellow in toluidine-blue stained sections (Figure 1J), the sporopollenin-containing exine can be stained with the dye Auramine-O. This staining is restricted to pollen grains, and to a thin band, of unknown origin, between the tapetum and the middle layer (Figure 1S). By contrast, in *gso1 gso2*, *cif3 cif4* and *tpst* mutants, yellow coloration and Auramine-O staining were also observed between the tapetum cells, extending to cell junctions with the middle layer (Figure 1L, T, Figure S4B, D, M, N). TEM analyses confirmed ectopic accumulation of electron-dense osmophilic material resembling sporopollenin both between and behind the tapetal cells of mutant anthers (Figure 1V and Figure S4O, P).

Tapetum cells become polarized as they differentiate allowing for directional secretion of sporopollenin precursors into the anther locule. (*5*). In order to address whether GSO1 signaling might influence tapetum polarization resulting in mislocalization of sporopollenin, we analyzed the expression and localization of the ABCG26 transporter involved in the secretion of pollen wall components from the tapetal cells. We found that a ABCG26:mTQ2 fusion protein expressed from the endogenous ABCG26 promoter was detected both inside the intracellular compartments and on the plasma membrane where it was polarly localized towards the locular matrix in wild-type plants (Figure S5C), as previously reported for the rice ABCG26 homologue (*16*). ABCG26:mTQ2 plasma membrane polarity was established at the early free microspore stage (stage 8; Figure S5A-C). In contrast, in the *gso1 gso2*, *cif3 cif4* and *tpst* mutants ABCG26:mTQ2 plasma membrane signal was detected predominantly between tapetal cells indicating a lack of cell polarization (Figure S5D-M). No change in expression was observed for the sporopollenin biosynthesis genes (Figure S6), suggesting that the accumulation of pollen wall material in the mutants is likely not due to enhanced sporopollenin biosynthesis, but may be due to polarity defects.

In summary, GSO1/2, CIF3/4 and TPST all appear to be necessary for normal tapetum function. Tapetum function in turn is necessary for normal pollen grain spacing within the locular fluid and for the targeted and regular deposition of pollen wall components onto the surface of developing pollen grains, suggesting that GSO-mediated signaling is necessary for the coordination of these processes.

### GSO receptors and are produced in the middle layer while CIF3 and CIF4 are expressed and sulfated in the tapetum cells

To understand the organization of the GSO signaling pathway in developing anthers, we studied the expression of pathway components. Transcriptional reporters revealed expression of *GSO1* and *GSO2* in the middle layer (Figure 2 A-B and Figure S7). Expression was initiated prior to meiosis of the pollen mother cell, at around stage 5 (Figure S7), significantly earlier than pollen wall deposition, and was maintained until the degradation of the middle layer at stage 11 (*13*). Translational reporter constructs revealed the presence of GSO1 and GSO2 proteins at the periphery of middle layer cells, consistent with plasma membrane localization (Figure 2 D-E and Figure S7). The GSO1 translational reporter fully complemented the pollen and tapetum phenotypes of *gso1 gso2* mutants (Figure S1B), confirming that this expression pattern accurately reflects native *GSO1* expression in the anther. Although both transcriptional reporters and fusion proteins were occasionally seen in the endothecium, neither gene expression nor protein accumulation was ever detected in the tapetum. Protein accumulation appeared uniform within the membrane of middle layer cells, similar to the situation previously reported in the embryo epidermis (*9*).

**Fig. 2.**
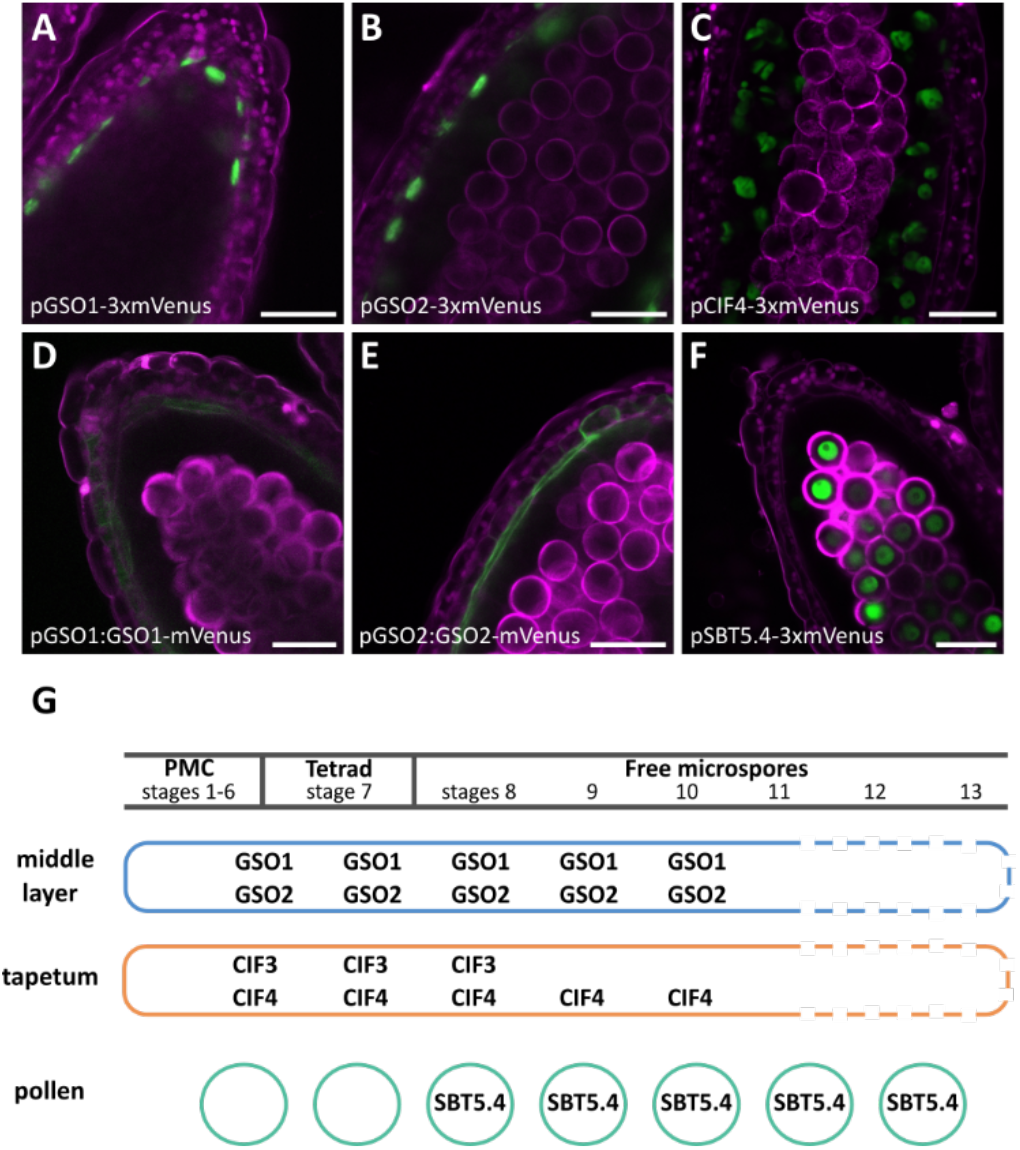
The GSO signaling pathway involves three different cell types. **(A-B)** Expression of the *GSO1* and *GSO2* genes in anthers using *pGSO1-3xmVenus* and *pGSO2-3xmVenus* reporter lines. **(C)** Expression of *CIF4* in anthers using *pCIF4-3xmVenus* reporter line. **(D-E)** Protein localization of the GSO1 and GSO2 receptor-like kinases in anthers using *pGSO1:GSO1-mVenus* and *pGSO2:GSO2-mVenus* reporter lines. **(F)** Expression of *SBT5.4* in anthers using *pSBT5.4-3xmVenus* reporter line. **(G)** Schematic illustration of the expression of GSO pathway components in anther cell types during development. Scale bars: A-F 20 μm.

In contrast, transcriptional reporters pCIF3-3xmVenus and pCIF4-3xmVenus revealed that the expression of *CIF3* and *CIF4* is restricted to the tapetum (Figure 2C, Figure S8) and starts at around the onset of PMC meiosis. While *CIF3* expression diminishes shortly after microspore release, CIF4 expression continues until tapetum degradation at stage 11 (Figure S8). Expression of the *CIF4* ORF either under its own promoter, or under the tapetum-specific *AMS* (*17*) (Figure S9), and *SHT* (*18*)(Figure S9) promoters, fully complemented the *cif3 cif4* phenotype (Figure S1G, H, J, O, Q) confirming that these expression patterns likely reflect the native expression of *CIF3* and *CIF4*.

To explore the spatial requirement for TPST, which acts cell autonomously in the Golgi apparatus during peptide secretion (*19*), we expressed the *TPST* ORF under the tapetum-specific *AMS* promoter in the *tpst-1* mutant. All transformed lines showed a complete or nearly complete complementation of the pollen and tapetum phenotypes of *tpst-1*, consistent with TPST activity being required for the production of sulfated peptide ligands in the tapetum (Figure S1L, R).

In summary, our results show that normal pollen wall deposition and tapetum function depend on the activity of CIF3 and CIF4 peptides sulfated and secreted from the tapetum, and their cognate receptors, GSO1 and GSO2 in the anther middle layer.

### A pollen specific subtilisin serine-protease, SBT5.4, can cleave the extended C-terminus of the CIF4 precursor

Analysis of the embryonic cuticle integrity signaling in seeds revealed a bidirectional signaling pathway in which an inactive, sulfated precursor of the TWS1 peptide requires C-terminal subtilase-mediated processing to release the mature and bioactive TWS1 peptide as a ligand for GSO1/2 (*9*). Like TWS1, both CIF3 and CIF4 possess C-terminal extensions (Figure 3A) suggesting that they may also require C-terminal processing for activation. We found that both the full length *CIF4* ORF, and a truncated version lacking the sequence encoding the part of the precursor C-terminal to the predicted active peptide, could complement the *cif3 cif4* mutant phenotype when expressed under the *CIF4* promoter or under the tapetum-specific *SHT* promoter in developing anthers (Figure S1G, I, J, K, O, P). The C-terminal extension of CIF4 is thus not required for activity. Our finding that C-terminal extensions in CIF-class ligands impair receptor binding (*9*), and the apparent contribution of the free C-terminus of CIF peptides to receptor/co-receptor complex formation (*11*), suggest that the C-terminal extension needs to be removed for bioactivity. We therefore tested the hypothesis that, as in the seed, subtilases could be involved in activating CIF peptides in the anther. We found that *SBT5.4*, one of numerous subtilase genes expressed in developing anthers (including *SBTs 1.8*, *3.7*, *3.12*, *3.16*, *3.18*, *4.13*, *4.15*, *5.1*, *5.4-5.6*), showed strong and apparently specific pollen-localized expression from stage 8, when the pollen wall starts to be laid down, until pollen maturity (Figure 2F, Figures S10, S11). SBT5.4 is thus a good candidate for CIF activation.

**Fig. 3.**
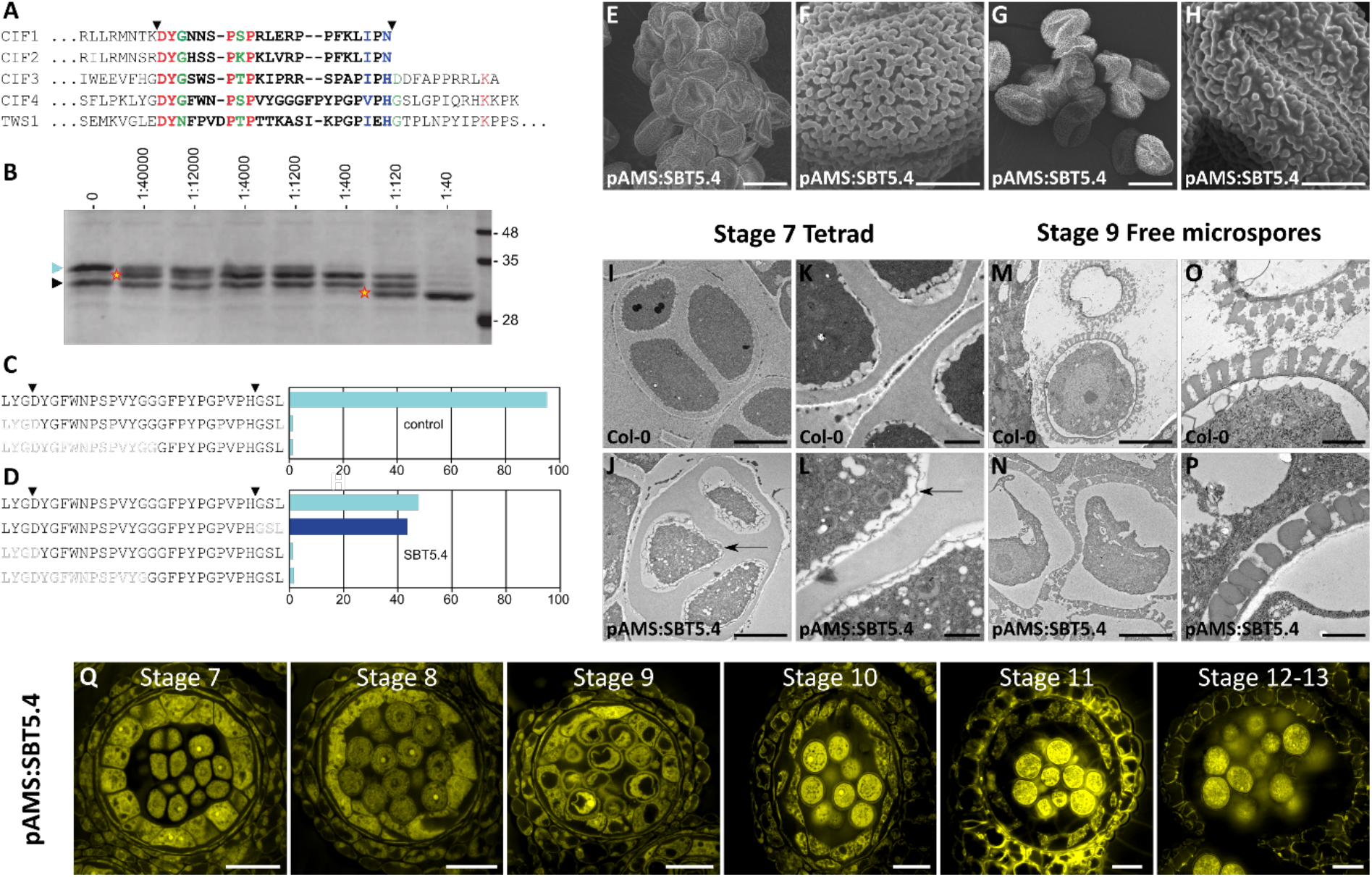
SBT5.4 can process the CIF4 precursor, and ectopic *SBT5.4* expression leads to deregulated pollen wall formation. **(A)** Sequence alignment of the C-termini of CIF family precursors, mature peptide sequences in bold face. Fully, strongly and weakly conserved residues (Clustal W; Gonnet Pam250 matrix) are highlighted in red, blue and green, respectively. Arrow heads indicate processing sites. **(B)** Coomassie-stained SDS-PAGE showing digests of recombinant GST-CIF4 (the full length and a truncated version of the precursor are marked by blue and black arrow heads, respectively) with SBT5.4 purified from tobacco leaves. The molar subtilase:substrate ratio is indicated for each lane; cleavage products highlighted by asterisks. **(C, D)** Cleavage of a synthetic CIF4 precursor peptide by SBT5.4. Bar graphs show relative abundance of the substrate peptide (top row) and fragments thereof for the control (C) and SBT5.4 (D) digests as the percentage of all peptides detected by MS/MS analysis. Arrow heads mark N- and C-terminal processing sites flanking the mature CIF4 peptide. The cleavage product generated by SBT5.4-specific processing of the C-terminal extension is highlighted in dark blue. **(E-H)** Scanning electron microscopy (SEM) images of *pAMS:SBT5.4* pollen from two independent lines in (E, F) and (G, H). Similar phenotypes were observed in 4 independent lines. **(I-P)** Transmission electron microscopy (TEM) of the pollen wall formation in the wild-type (I, K, M, O) and the *pAMS:SBT5.4* (J, L, N, P) anthers at the tetrad stage (stage 7) and the free microspore stage (stage 9). Black arrows indicate putative detached primexine in the *pAMS:SBT5.4* tetrads. **(Q)** Pollen and anther development in the *pAMS:SBT5.4* line at different developmental stages. Scale bars: E, G, Q 20 μm; F, H, I-J, M-N O 5 μm; K-L, O-P 1 μm.

*SBT5.4* loss-of-function mutants did not show any defects in pollen development (Figure S12), possibly due to the presence of functionally redundant proteases. Furthermore, a triple mutant lacking function of *SBT5.4*, *5.5* and *5*.6, all of which showed similar expression patterns in the developing pollen, also failed to show any pollen phenotype (Figure S11, 12).

We therefore tested directly whether SBT5.4 is able to cleave the CIF4 precursor. N-terminally GST-tagged proCIF4 expressed in *E. coli* was co-incubated with SBT5.4:(His)6 transiently expressed in tobacco (*N. benthamiana*) leaves and purified from cell wall extracts. Several cleavage products were generated upon co-incubation with SBT5.4, but not with extracts from untransformed plants, indicating that proCIF4 is processed by SBT5.4 (Figure 3B). To confirm cleavage at sites relevant for CIF4 maturation, a synthetic CIF4 peptide extended by three amino acids of the precursor at either end was used as substrate for recombinant SBT5.4. Mass spectrometry analysis of cleavage products revealed specific cleavage only at the C-terminal processing site between His89 and Gly90 (Figure 3C, D). SBT5.4, a subtilase specifically expressed in pollen grains from stage 8 onwards, is thus able to remove the C-terminal extension for CIF4.

To further confirm the potential role of SBT5.4 we expressed it under the *AMS* promoter which drives expression specifically in the tapetum, from the point of PMC meiosis onwards (Figure S9) (and thus earlier than the initiation of endogenous *SBT5.4* expression in pollen grains). This led to strong pollen phenotypes (Figure 3). Pollen in these lines tended to fuse together in a mass (Fig. 3E-H). However, defects in these lines were very distinct from those observed in *gso1 gso2* or *cif3 cif4* mutants. Firstly, tapetum hypertrophy was either weaker or absent (Figure 3Q). Secondly, and again unlike the situation in loss of function *gso1 gso2* or *cif3 cif4* mutants where the reticulate patterning of the pollen wall is still visible in most peripheral pollen grains (Figure 1C-F), the patterning of the pollen wall was severely compromised in lines expressing the *pAMS:SBT5.4* construct (Figure 3F, H). Defects varied from loss/fusion of tecta, giving rise to a “broken” appearance, to an apparent massive de-regulation of pollen wall deposition, giving pollen grains a “lumpy” appearance (Figure 3F, H). Deregulation of pollen wall deposition could be observed as early as the tetrad stage in the *pAMS-SBT5.4* lines. At this stage the organization of the primexine, which serves as a scaffold for the pollen wall formation, was abnormal (Figure 3I-L). At later stages, the bacula appeared to form very close to each other resulting in abnormally dense exine (Figure 3 N, P). Bacula also appeared to be poorly attachment to the pollen surface (Figure 3P). Thus, the ectopic expression of the SBT5.4 leads to strong deregulation of the pollen wall formation.

Our results strongly support the hypothesis that the strict spatial and temporal regulation of CIF-cleaving SBT activity in developing pollen grains is a critical factor in ensuring the organized deposition of the pollen wall.

### GSO1 expression in the tapetum interferes with pathway function

Our results suggest a model in which tapetum-derived peptides, processed by pollen-derived subtilases, must diffuse to the middle layer to activate GSO-mediated signalling (Figure 4G). To test this model further, we expressed full-length GSO1 and a truncated version lacking the cytoplasmic kinase domain (GSO1 ΔKinase) under the tapetum-specific *AMS* promoter. ΔKinase versions of LRR-RLKs have previously been shown to provoke dominant negative phenotypes, possibly through ligand sequestration (*20*). Consistent with this, wild-type plants transformed with *pAMS:GSO1-ΔKinase* showed a strong loss of function phenotype identical to the *gso1 gso2* mutant (tapetum hypertrophy with both ectopic and defective pollen wall deposition) (Figure 4A-C). This could be explained by the sequestration of activated CIF peptides, which according to our model need to diffuse around tapetum cells to reach their receptors in the middle layer. In contrast, wild-type plants transformed with *pAMS:GSO1* presented a different phenotype with disrupted pollen wall patterning, frequent pollen death, pollen grain adhesions, but no tapetum hypertrophy (Figure 4D-F). These phenotypes are reminiscent of those observed upon *SBT5.4* expression under the *AMS* promoter, suggesting that ectopic GSO1 expression in the tapetum may cause a deregulation of pathway activation.

**Fig. 4.**
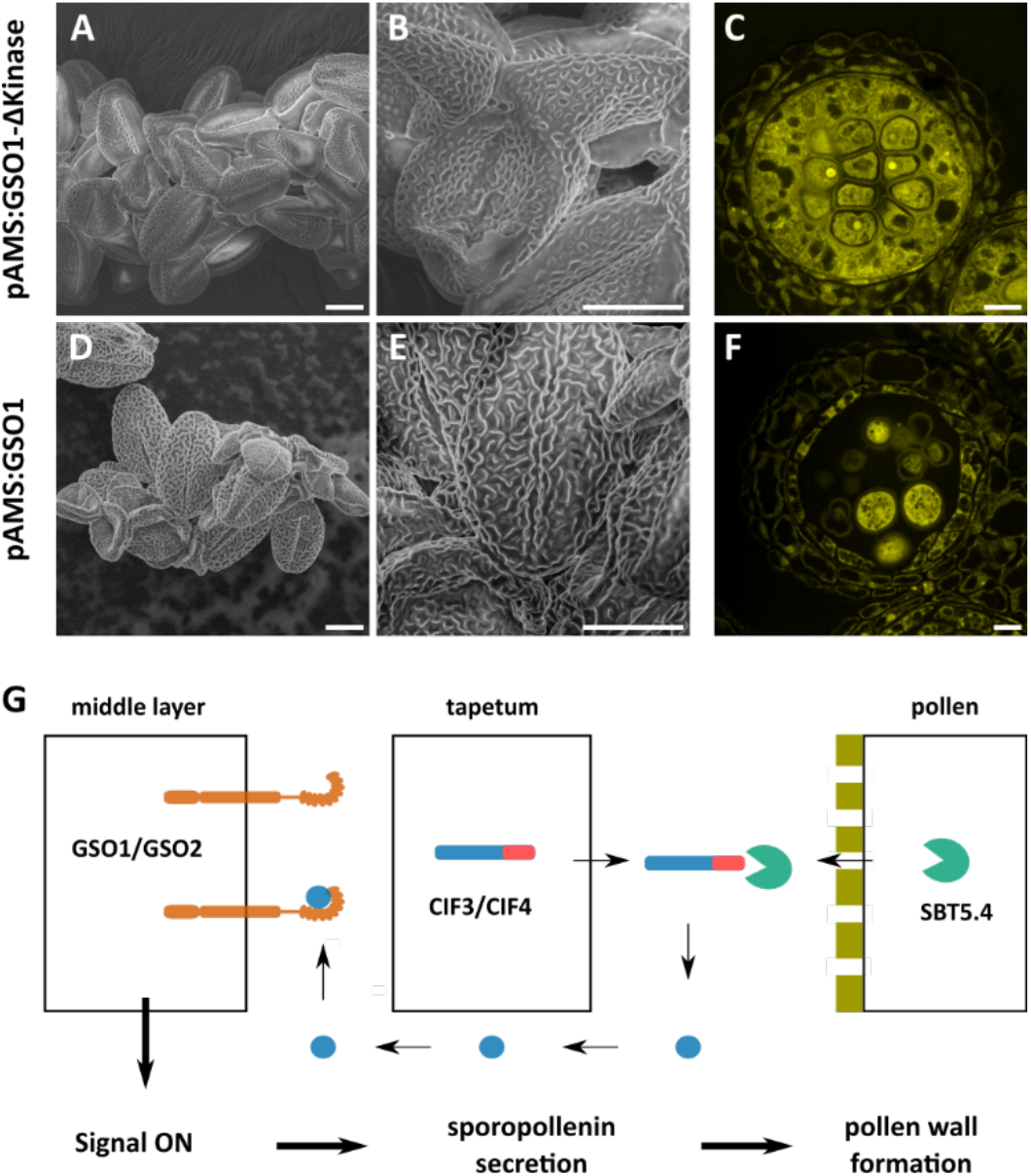
The complex spatial organization of GSO pathway components is necessary for controlled tapetum development. **(A-C)** Scanning electron microscopy (SEM) images and histological cross-section of anthers of the *pAMS:GSO1-ΔKinase* line. Phenotype confirmed in 7 independent lines. **(D-F)** Scanning electron microscopy (SEM) images and histological cross-section of anthers of the *pAMS:GSO1* line. Phenotype confirmed in 5 independent lines. **(G)** Taken together our data support the following model for the coordination of pollen wall formation with tapetum development. The CIF precursors produced in the tapetum are processed by the pollen-derived subtilases. The activated peptides diffuse between tapetum cells to bind GSO receptors in the middle layer which induces downstream signaling leading to polarized sporopollenin secretion towards the pollen grains. The completion of the pollen wall on the pollen surface prevents the interaction of subtilases with their substrates, the uncleaved peptides are inactive and this attenuating signaling. Scale bars: 10 μm.

## Discussion

The coordination of tapetum activity (polar secretion of nutrients, enzymes, pollen wall components, and the locular matrix) with pollen development (growth, and initial patterning of the pollen wall) is indispensable for the generation of viable pollen grains. Taken together our data indicate that this coordination involves a molecular dialogue between the tapetum, the developing pollen grains and the middle layer. The localization of GSO1 and GSO2 receptors on the membrane of middle-layer cells, and expression of CIF3 and CIF4 precursor-encoding genes in the tapetum, initiate significantly before microspore release from tetrads and the onset of pollen wall deposition (Figure 2G, S7 and S8). However, our phenotypic data show that this early production of signaling pathway components, which is also observed in the seed system (*9*), may “poise” the signaling system for action, but is not necessary for early stages of tapetum or pollen grain development, since loss of function phenotypes in both ligand and receptor mutant backgrounds only appear after microspore release.

In contrast, our results suggest that the trigger in the pathway is the production of pollen-derived CIF-activating proteases including SBT5.4, the microspore-specific expression of which is synchronized perfectly with the onset of pollen wall deposition (Figure 2G, S11). The importance of accurately timing pathway activation is underlined by the dramatic effects of mis-expressing SBT5.4 early in the tapetum, which would be predicted to trigger a precocious activation of GSO-mediated signalling through constitutive ligand activation. The highly disrupted deposition of pollen-wall components observed in these lines, including the difficulty of wall components to anchor to the pollen surface could be a consequence of a lack of coordination between the production of pollen wall component by the tapetum, and microspore release from tetrads.

Our model predicts that subtilases produced in immature pollen grains mediate the timely processing and liberation of activated CIF3/4 peptides in the pollen wall or locular matrix. Activated peptides must then diffuse between tapetum cells to the middle layer where they can bind to GSO1 and GSO2, ultimately leading to the activation of the tapetum, and the organized secretion of pollen wall components into the anther locule. Pollen wall completion will attenuate signaling activity by separating subtilases from their substrates, thus adjusting tapetum activity to the needs of the pollen grains (Figure 4G).

This model raises the intriguing question of why GSO1 and GSO2 are expressed in the middle layer, rather than in the tapetum. One possible explanation is that separating pro-peptide and receptor production prevents untimely receptor-ligand interaction in the secretory pathway. However our published data suggest that C-terminally extended CIF peptides show impaired receptor binding (*9*) and, consistent with this, co-expression of the TWS1 precursor and the GSO receptors in the embryo epidermis does not lead to receptor activation. A second possibility, supported by out data (Figure 4D-F), is that GSO signaling interferes with other signaling pathways if it occurs in the tapetum, for example through competition for SERK co-receptors which are known to interact with GSO1 and GSO2 but also with other receptors involved in anther development (*11*, *21*). A final possibility is that the directional output of GSO signaling from the middle layer helps to polarize secretion from the tapetum, as suggested by the ectopic deposition of sporopollenin in pathway mutants.

In conclusion, our work has revealed that a complex peptide-mediated dialogue spanning three distinct cell types is necessary to synchronize secretion and deposition of pollen wall components within the anther locule. The middle layer, an understudied and poorly understood anther tissue, plays an important role in this dialogue, apparently acting as an organizing hub for the activation and associated polarization of the neighboring tapetum.

## Materials and Methods

### Plant material and growth conditions

Seeds were sown on half-strength MS (Murashige Skoog) medium with 1% sucrose and 1% agar, stratified for 2 days at 4°C and grown for 7 days in short-day conditions (8h light / 16h dark). Seedlings were then transferred to soil and grown for 3 more weeks in short-day conditions (8h light / 16h dark) and subsequently transferred to long-day conditions (16h light / 8h dark) to promote flowering. Some of the transgenic lines described were examined in the T1 generation; these were first selected on half-strength MS medium with 1% sucrose and 1% agar supplemented with either 50 μg/mL kanamycin or 10 μg/ml glufosinate ammonium (Basta).

For *promoter::GUS(uidA)* reporter lines, seeds were incubated over night at −75 °C, re-suspended in 0.1 % agar, stratified for 48 hours at 4°C in the dark, and sown on either half-strength MS (Murashige Skoog), 1% sucrose, 0.6 % agar plates, or a mix of potting compost with 3.6 % (v/v) sand and 7.2 % (v/v) perlite. Plants were grown under short-day conditions (12 h photoperiod) at 22 °C and 100-120 μE white light.

Mutant alleles used were *gso1-1* (SALK_064029) *gso2-1* (SALK_130637) (*9*, *22*), *gso1-2* (SALK_103965) *gso2-2* (SALK_143123) (*10*), *cif3-2* (GABI_516E10) *cif4-1* (CRISPR mutant line) (*11*), *tpst-1* (SALK_009847) (*19*), *sbt5.4* (*At5g59810*, SALK_025087), *sbt5.5* (*At5g45640*, SALK_107233), *sbt5.6* (*At5g45650*, CRISPR mutant generated as described below) (Fig. S12); genotyping primers used are listed in Suppl. Table 1. The *pGSO1:GSO1-mVenus* and *pGSO2:GSO2-mVenus* lines are described in (*9*).

### Generation of transgenic plant lines

Gateway technology (Invitrogen^™^) was used for the production of genetic constructs. For the 3xmVenus transcriptional reporter lines, the following promoter fragments were amplified by PCR from Arabidopsis (Col-0) genomic DNA: *pSBT5.4* −2639 bp to −1 bp, *pSBT5.5* −1903 bp to −1 bp, *pSBT5.6* −4144 bp to −1 bp, *pAMS* −2618 bp to −1 bp, *pSHT* −859 bp to −1 bp, *pGSO1* −5583 bp to −1 bp, *pGSO2* −3895 bp to −1 bp, *pCIF3* −2092 bp to −1 bp, *pCIF4* −2201 bp to −1 bp. The fragments were inserted into pDONR P4-P1R and recombined with *3xmVenus-N7* pDONR211, *OCS* terminator pDONR P2R-P3 (containing STOP codon followed by the *OCTOPINE SYNTHASE* terminator) and pK7m34GW destination vector (with kanamycin *in planta* resistance); this yielded plasmids containing *pSBT5.4-3xmVenus*, *pSBT5.5-3xmVenus*, *pSBT5.6-3xmVenus*, *pAMS-3xmVenus*, *pSHT-3xmVenus*. To construct *pGSO1-3xmVenus*, *pGSO2-3xmVenus*, *pCIF3-3xmVenus*, and *pCIF4-3xmVenus*, promoter regions cloned into pDONR P4-P1R vectors and *NLS-3xmVenus* in pDONRZ were integrated into the pB7m24GW,3 destination vector (with Basta *in planta* resistance and with the *35S* terminator) by LR reaction. These constructs were transformed into the Col-0 background.

For the *SBT5.4* promoter::GUS(uidA) reporter line 1592 bp upstream of the ATG translational start were amplified from Arabidopsis (Col-0) genomic DNA and cloned into the *NotI/SalI* sites of pGreen 0029 (*23*) containing a promoterless *β-glucuronidase (GUS):green fluorescent protein (GFP)* fusion reporter gene from pCambia 1303 (CAMBIA GPO Box 3200 Canberra, ACT 2601 Australia) followed by the 282 bp *OCS* terminator.

To produce *GSO1* mis-expression constructs, the *GSO1* genomic fragment lacking the cytoplasmic kinase-encoding domain (from the ATG (+1 bp) to +2850 bp) was inserted into pDONR211 and recombined with *pAMS* pDONR P4-P1R (see above), *OCS* terminator pDONR P2R-P3 (see above) and pK7m34GW destination vector to yield the *pAMS:GSO1-ΔKinase* construct. Similarly, the entire *GSO1* genomic fragment from ATG (+1 bp) to the STOP codon (+3826 bp) was inserted into pDONR211 and recombined with *pAMS* pDONR P4-P1R (see above), *OCS* terminator pDONR P2R-P3 and pK7m34GW destination vectors to yield the *pAMS:GSO1* vector. These constructs were transformed into Col-0 background.

The *CIF4* complementation construct was produced by amplifying the *CIF4* genomic fragment including the promoter region and the entire coding sequence from −4242 bp to + 309 bp (including the STOP codon) and introducing it into pDONR211. Alternatively, a truncated version of the *CIF4* genomic fragment from −4242 bp to +267 bp with an artificial STOP codon before the C-terminal extension was introduced into pDONR211. The *CIF4* terminator fragment from +310 bp to + 2541bp was inserted into pDONR P2R-P3. Genomic fragments were recombined into the pK7m34GW destination vector to yield *pCIF4:CIF4* or *pCIF4:CIF4_truncated* constructs which were then transformed into the *cif3-2 cif4-1* double mutant.

To produce *CIF4* expression constructs, *CIF4* coding sequence from +1 bp (ATG) to +309 bp (STOP codon) was introduced into pDONR211. The construct was recombined with either *pAMS* pDONR P4-P1R or *pSHT* pDONR P4-P1R and with *OCS* terminator pDONR P2R-P3 and pK7m34GW destination vectors to yield the *pAMS:CIF4* and *pSHT:CIF4* constructs. Alternatively, a truncated version of the *CIF4* coding sequence including N-terminal part from +1 bp (ATG) to +267 bp (and therefore lacking the C-terminus that is removed in the mature peptide) was introduced into pDONR211. The construct was recombined with *pSHT* pDONR P4-P1R and with *OCS* terminator pDONR P2R-P3 and pK7m34GW destination vectors to yield the *pSHT:CIF4_ truncated* construct. These constructs were transformed into the *cif3-2 cif4-1* double mutant.

To produce the *TPST* complementation construct, a plasmid containing the *TPST* ORF in pDONR211 (*9*) was recombined with the *pAMS* pDONR P4-P1R, OCS terminator pDONR P2R-P3 and the pB7m34GW destination vector to yield the *pAMS:TPST* construct. This construct was introduced into the *tpst-1* background.

For ectopic expression of SBT5.4 in the tapetum, the *SBT5.4* coding sequence from +1 bp (ATG) to +2337 bp (STOP codon) was introduced into pDONR211. The construct was recombined with *pAMS* pDONR P4-P1R and with OCS terminator pDONR P2R-P3 and pK7m34GW destination vectors to yield the *pAMS:SBT5.4* construct. This construct was transformed into the Col-0 background.

For the generation of the CRISPR allele of *SBT5.6* the method described by Stuttmann and co-workers was used (*24*). Two guides cutting at the beginning of the gene coding sequence were designed and inserted into the pDGE332 and pDGE334 shuttle vectors. The two shuttle vectors were then recombined into the pDGE347 recipient vector containing FastRed selection marker. The construct was transformed into the *sbt5.4 sbt5.5* double mutant background. The transgenic plants were selected with FastRed and analyzed for the presence of the mutation. The CRISPR allele was shown to contain a 104 bp deletion in the second exon of the *SBT5.6* gene.

To generate the *pABCG26:ABCG26-mTQ2* translational reporter line, the *ABCG26* genomic fragment including the promoter (−583 bp to −1 bp) and the coding sequence with introns up to but not including the STOP codon was amplified by PCR from Arabidopsis (Col-0) genomic DNA and cloned into the pENTR 5’-TOPO vector. The *mTQ2* ORF with four alanine codons for a small N-terminal linker replacing the start codon was cloned into the pDONR211 vector. The *pABCG26:ABCG26* pENTR 5’-TOPO, the *mTQ2* pDONR211 vector were recombined with the *OCS* terminator pDONR P2R-P3 vector and pB7m34GW destination vector. The construct was transformed into the Col-0, *gso1-1 gso2-1, cif3-2 cif4-1* and *tpst-1* backgrounds.

For plant transformation, the constructs were first introduced into *Agrobacterium tumefaciens* strains C58pMP90 by electroporation, and then transformed into Col-0 plants or the respective mutant genotypes by floral dip as described in (*25*).

Primers used are listed in Suppl. Table 1.

### Histology

Inflorescences were fixed with FAA (50% (v/v) ethanol, 5% (v/v) acetic acid, 3.7% (v/v) formaldehyde) overnight, dehydrated in a graded series of 50%, 60%, 70%, 85%, 95% and 100% ethanol for 1 h each, then further incubated overnight in fresh 100% ethanol. The samples were then incubated in 50% ethanol/50% Technovit 7100 base liquid (v/v) for 4h and then in 25% ethanol/75% Technovit 7100 base liquid (v/v) overnight. The samples were infiltrated in Technovit 7100 infiltration solution (1g hardener I in 100 ml Technovit 7100 base liquid) with vacuum for 2h and further incubated for 6 days. All steps above were conducted at room temperature (RT) with gentle agitation. The samples were polymerized with Technovit 7100 polymerisation solution (100 μl Technovit 7100 hardener II in 1,5 ml infiltration solution) at RT for 6 hours. Transverse section of 3μm were cut using a Leica Microtome HM355S.

For histological analysis, the sections were stained with 0.01% (w/v) acriflavine in H2O for 5 min, mounted in Vectashield (Vector Laboratories) and observed using TCS SP5 or TCS SP8 confocal microscopes (Leica) with excitation at 488 nm and emission at 492-551 nm.

Alternatively, the sections were stained with the 0.05 % (w/v) Toluidine blue in H_2_O for 1 minute, mounted in Entellan mounting medium (Sigma) and observed under Zeiss Axio Imager M2 microscope.

For histochemical staining of GUS activity (*26*) in the *promoter::GUS(uidA)* reporter lines, plants were vacuum-infiltrated with 2 mM each of potassium ferrocyanide, potassium ferricyanide and 5-bromo-4-chloro-3-indolyl-β-D-glucuronic acid (X-Gluc, Biochemical DIRECT) in 50 mM sodium phosphate buffer pH 7.0 with 0.1% Triton X-100 (v/v) for 2 min at 80 mbar. After 24 h at 37°C, the tissue was destained in a graded ethanol series (20, 35, 50, 70, 80, 90% (v/v)). Pictures were taken with a Nikon D5000 digital camera and at a dissecting microscope (Stemi SV11; Carl Zeiss Microscopy, Jena, Germany) using a SPOT RT KE Color Mosaic Camera model 7.2 with SPOT software (Visitron Systems; Puchheim, Germany).

### Clearsee tissue clearing

Inflorescences were fixed in 4% paraformaldehyde in PBS at 4°C under vacuum for 2h and subsequently incubated overnight at 4°C. The samples were washed twice with PBS and cleared with Clearsee Alpha solution (*27*) (10% (w/v) xylitol powder, 15% (w/v) sodium deoxycholate, 25% (w/v) urea and 0.63% (w/v) sodium sulfite) for 1 week changing to a fresh solution every 2 days at RT.

After clearing, the samples were incubated in 0.1% (w/v) Auramine O dissolved in Clearsee Alpha overnight, washed 3 times for 20 min each with Clearsee Alpha solution. The anthers were dissected from the inflorescences, mounted in Clearsee Alpha solution and observed using a confocal TCS SP5 or TCS SP8 confocal microscope (Leica). Auramine O was excited at 488 nm and the emission was collected at 500-570 nm.

### Scanning and cryo-scanning electron microscopy

Pollen grains were observed in the scanning electron microscope SEM FEG FEI Quanta 250 at 5kV voltage or, alternatively, using SEM HIROX 3000 at 10 kV.

For cryo-SEM, anthers were immobilized on a specimen mount coated with modeling clay and rapidly frozen in liquid nitrogen for at least 15 s under vacuum. The specimen was transported under vacuum into the cryo-SEM MEB FEG FEI Quanta 250 precooled to −140°C. The frozen anthers were fractured at −140°C, then sublimed at −90°C to remove ice accumulations, immediately refrozen to −140°C and coated with gold. The anthers were observed at 5 kV.

### Transmission electron microscopy

Flower buds at appropriate developmental stages were fixed with 4 % (w/v) formaldehyde and 2 % (w/v) glutaraldehyde in 0.1 M phosphate buffer (pH 7.2) (PB) under vacuum (0.6 bar) at 4°C for 1h during which the vacuum was slowly broken 3 times, then incubated in fresh fixative solution at 4°C overnight. The samples were washed three times in PB, postfixed for 2h in 1 % (w/v) osmium tetroxide in PB at room temperature (RT), rinsed 5 times for 5 min in PB and dehydrated under vacuum in a graded ethanol series (30 to 100% v/v) with 20 min incubations in each bath at RT. The samples were then infiltrated with low viscosity Spurr resin (EMS Catalog #14300) using a graded series in ethanol (33%, 66% and twice 100% v/v Spurr resin) at 4°C with each incubation lasting 24h (including 20min under vacuum). The samples were polymerized in fresh Spurr resin at 60°C for 18h. Ultrathin sections (70 nm) were prepared using UC7 Leica Ultramicrotome, placed on formvar-coated grids, then poststained with 2% uranyl acetate and lead citrate. Sections were examined under JEOL 1400 transmission electron microscope at 120kV and imaged with the Gatan Rio 16 camera.

### Confocal microscopy

For the mVenus-expressing transcriptional and translational reporter lines, anthers were stained with 20 μg/ml propidium iodide solution and examined using TCS SP5 (Leica) or ZEISS 710 confocal microscopes with excitation at 514 nm and emission at 526-560 nm for mVenus, and 605-745 nm for propidium iodide.

For the *pABCG26:ABCG26-mTQ2* reporter lines, the anthers were stained with 20 μg/ml propidium iodide solution and examined using ZEISS 710 confocal microscopes with sequential excitation at 458 nm for mTQ2 and emission peak at 490 nm, as well as with excitation at 514 nm and emission peak at 642 nm for propidium iodide. 40 anthers per genotype from the same developmental stage were imaged.

### Peptide cleavage assay and cleavage product analysis

The *SBT5.4* ORF was amplified by PCR from a RIKEN full-length cDNA clone (RAFL19-27-C18). Primers included *Kpn*I and *Xba*I restriction sites and six terminal His codons in the reverse primer. The PCR product was cloned between the CaMV 35S promoter and terminator in pART7 (*28*). The expression cassette was then transferred into the *Not*I site of the binary vector pART27, and introduced into *Agrobacterium tumefaciens* (C58C1). C-terminally His-tagged SBT5.4 was transiently expressed in *N. benthamiana* by agro-infiltration as described (*29*). The leaf apoplast was extracted four to five days after infiltration with 50 mM Na_2_HPO_4_/NaH_2_PO_4_ pH 7.0, 200 mM KCl, 1 μM pepstatin, 10 μM E64. Extracts were subjected to affinity chromatography on 500 μl Ni-NTA agarose slurry according to protocols provided by the manufacturer (Qiagen; Hilden, Germany).

The *CIF4* ORF, without the sequence coding for the signal peptide, was amplified by PCR from cDNA using primers that included a *Bam*HI site at the 5’-end, and six His codons and an *Eco*RI site at the 3’-end. The PCR product was cloned downstream and in frame with the glutathione *S*-transferase (GST) ORF in pGEX-3X (GE Healthcare). The expression construct was transfected into *E. coli* BL21-CodonPlus(DE3)-RIL (Agilent Technologies). Cells were grown in 250 ml LB medium at 37 °C and 200 rpm, until OD_600_ reached 0.6 to 0.8. Subsequently, the culture was cooled down to 30 °C and after 20 min at 200 rpm, 1 mM IPTG was added to induce protein expression. Cells were harvested by centrifugation after 2 hrs, and lysed by sonication in 50 mM Na_2_HPO_4_/NaH_2_PO_4_ pH 7.0, 150 mM NaCl, 1 mM PMSF, 5 mM benzamidine hydrochloride, and a spatula tip DNAseI. Four mM imidazole was added to the cleared lysate, and the recombinant protein was purified on Ni-NTA agarose following the manfacturer’s instructions (Qiagen). The eluate (3x 500 μl) was concentrated by ultrafiltration (Vivaspin concentrators, 10 kDa molecular weight cut-off; Sartorius; Göttingen, Germany) and dialyzed against 50 mM Na_2_HPO_4_/NaH_2_PO_4_ pH 6.0, 300 mM NaCl.

Two and a half μg GST-CIF4-His in a total volume of 10 μl 50 mM Na_2_HPO_4_/NaH_2_PO_4_ pH 6.0, 300 mM NaCl were incubated at 25°C with recombinant SBT5.4 at the indicated concentrations. As negative control, apoplastic washes from *N. benthamiana* plants that had been agro-infiltrated with the empty pART27 vector were subjected to a mock-purification, and an equal volume of the final fraction was used in the assay. The reaction was stopped after 2 hrs by addition of 4x SDS-PAGE sample buffer. The digest was analyzed by SDS-PAGE (15%) followed by Coomassie-Brilliant Blue (Instant Blue; Abcam, Cambridge, UK) staining.

For mass spectrometric analysis of the cleavage site, 6 μg of a synthetic CIF4 peptide extended at both the N- and the C-terminus by three precursor-derived amino acid residues were digested with 50 ng recombinant SBT5.4 and a mock-purified fraction as a control. The reaction was stopped after 1 hour at 25°C by addition of 1% trifluoroacetic acid. Reaction products were purified on C18 stage tips equilibrated in 0.5% (v/v) acetic acid as described (*30*). Peptides were eluted in 0.5% (v/v) acetic acid, 50% (v/v) acetonitrile, evaporated to dryness in a SpeedVac concentrator (Savant Instruments; Holbrook, NY), and stored at −20°C until further analysis. Digests were analyzed by LC-MS/MS using an UltiMate 3000 RSLCnano system (Dionex, Thermo Fisher Scientific) coupled to an Orbitrap mass spectrometer (Exploris 480, Thermo Fisher Scientific). The Exploris 480 was operated under XCalibur software (version 4.4, Thermo Fisher Scientific) and calibrated internally (*31*). Peptides were trapped on a pre-column (C18 PepMap100, 5μm x 5mm, Thermo Fisher Scientific) and then eluted from a 75 μm x 250 mm analytical C_18_ column (NanoEase, Waters) by a linear gradient from 2 to 55% acetonitrile over 30 min. MS spectra (m/z = 200-2000) were acquired at a resolution of 60,000 at m/z = 200. Data dependent MS/MS mass spectra were generated for the 30 most abundant peptide precursors using high energy collision dissociation (HCD) fragmentation at a resolution of 15000 and a normalized collision energy of 30. Peptides were identified by a Mascot 2.6 search (Matrix Science, UK) and quantified using peak areas calculated from extracted ion chromatograms in XCalibur 4.4 (Thermo Fischer Scientific).

### Pollen viability and germination assays, seed count

Pollen viability was tested by staining mature anthers and pollen with Alexander solution (*32*). For the pollen germination assays mature pollen grains were transferred onto medium containing 0.5% low melting agarose, 18% sucrose, 0.01% boric acid, 1 mM CaCl_2_, 1mM Ca(NO_3_)_2_, 1 mM KCl, 0.03% casein enzymatic hydrolysate, 0.01% ferric ammonium citrate, 0.01% myo-inositol and 0.25 mM spermidine, pH 8.0 for 8h at 25°C. The images were taken with Zeiss Axio Imager M2 microscope.

For the seed number analysis, seeds were counted in the 4^th^-8^th^ silique of the respective plants. 50 siliques per genotype were counted. Unpaired two-tailed t-tests were used for statistical analysis.

### qRT-PCR

Whole inflorescences containing unopened flowers (from the first closed flower upwards) were collected with 4 independent replicates per genotype. RNA was extracted using the Spectrum Plant Total RNA kit (Sigma-Aldrich) and the DNA was removed using the TURBO DNA-free kit (Invitrogen). The cDNA was generated using SuperScript IV VILO Master Mix (Thermo Fischer) with 500 ng RNA per sample. The cDNA was then diluted 1:50. The qRT–PCR was performed using Applied Biosystems Fast SYBR Green Master Mix. The PCR reaction was as follows: 95°C for 10 min, 50 cycles of 95°C for 15 s, and 60°C for 30 s. Expression of *PP2AA3* gene (*At1g13320*) was used as standard (*33*, *34*). Similar results were obtained when the *AP2M* gene (*AT5G46630*) was used as the standard (*33*, *35*). The PCR efficiency was calculated from the standard curve amplification data: *ACOS5* 108%, *PKSA* 111%, *PKSB* 93%, *MS2* 100%, *TKPR2* 86%, *CYP703A2* 101%, *CYP704B1* 83%, *PP2AA3* 97%. Expression levels are presented as E^-ΔCt^ with ΔCt = Ct_GOI_-Ct_PP2AA3_. Primers used are listed in Suppl. Table 1.

## Acknowledgments

We thank Audrey Creff, Alexis Lacroix, Patrice Bolland, Justin Berger, Isabelle Desbouchages and Hervé Leyral for technical assistance, Cornelia Schmitt for the cloning of expression constructs, Cindy Vial, Stéphanie Maurin, Laureen Grangier and Nelly Camilleri for administrative assistance. Mass spectrometry analyses were performed at the Core Facility Hohenheim, Mass Spectrometry Unit (University of Hohenheim, Stuttgart, Germany). SEM and TEM images were acquired at the Centre Technologique des Microstructures, Université Lyon1. We acknowledge the contribution of SFR Biosciences (UMS3444/CNRS, US8/Inserm, ENS de Lyon, UCBL) facilities: C. Lionet, E. Chatre, and J. Brocard at the LBI-PLATIM-MICROSCOPY for assistance with imaging.

## Author contributions

G.I. led the study. G.I., N.G. and A.Sc. obtained funding for the study. G.I., A.Sc., N.G., and A.St. supervised the work. J.T., S.Br., S.Bo., N.M.D. and S.F. carried out the experiments. All authors were involved in the analysis of the results. G.I., J.T. and A.Sc. wrote the paper with input from all authors.

## Data and materials availability

All lines used in the study will be provided upon signature of an appropriate Material Transfer Agreement. All data is available in the main text or the supplementary materials.

